# Why phylogenies compress so well: combinatorial guarantees under the Infinite Sites Model

**DOI:** 10.64898/2026.03.18.712055

**Authors:** Veronika Hendrychová, Karel Břinda

## Abstract

One important question in bacterial genomics is how to represent and search modern million-genome collections at scale. Phylogenetic compression effectively addresses this by guiding compression and search via evolutionary history, and many related methods similarly rely on tree- and ordering-based heuristics that leverage the same underlying phylogenetic signal. Yet, the mathematical principles underlying phylogenetic compression remain little understood. Here, we introduce the first formal framework to model phylogenetic compression mechanisms. We study genome collections represented as RLE-compressed SNP, *k*-mer, unitig, and uniq-row matrices and formulate compression as an optimization problem over genome orderings. We prove that while the problem is NP-hard for arbitrary data, for genomes following the Infinite Sites Model it becomes optimally solvable in polynomial time via Neighbor Joining (NJ). Finally, we experimentally validate the model’s predictions with real bacterial datasets using an exact Traveling Salesperson Problem (TSP). We demonstrate that, despite numerous simplifying assumptions, NJ orderings achieve near-optimal compression across dataset types, representations, and *k*-mer ranges. Altogether, these results explain the mathematical principles underlying the efficacy of phylogenetic compression and, more generally, the success of tree-based compression and indexing heuristics across bacterial genomics.

## 1 Introduction

The rapid growth of bacterial genome collections presents a major challenge for their compression and search. Modern collections contain millions of bacterial assemblies: AllTheBacteria 2.8 million [1], NCBI-asm 3.0 million, and GTDB [2] 0.7 million genomes. When complemented by metagenome-assembled genomes, the total amount of distinct publicly available bacterial genomes exceeds 10 million [3] and continues to grow rapidly. The compression, distribution, and search of collections at this scale present a major algorithmic challenge.

Phylogenetic compression [4] has emerged as a high-level strategy to scale both compression and search well beyond the million-genome regime. Implemented in MiniPhy [4], it works as follows: genomes are phylogenetically grouped into small batches of related genomes, for instance based on their species, and then each batch is reordered according to its estimated evolutionary history. In practice, this organizes a large collection into a series of many (usually hundreds) relatively small trees whose leaves are individual genomes. This seemingly simple process makes even very large collections substantially more amenable to downstream compression [5,6] and indexing [4,7] workflows, yielding space improvements of one to three orders of magnitude over prior protocols. Furthermore, the same principle extends across a broad range of representations and downstream data structures, including assemblies, de Bruijn graphs, *k*-mer indexes, and sketches.

Phylogeny-guided reordering is the key compression enabler: by placing closely related genomes adjacent to one another, phylogenetic ordering co-localizes shared underlying evolutionary redundancies that compressors and indexers can easily exploit. The utility of data reordering for boosting compression is well established across computer science, and numerous scalable genome indexing methodologies now implement some form of structure-aware heuristic [8,9,10,11]. In microbial genomics, these methods are particularly effective because they invariably leverage the same phylogenetic signal.

However, despite these empirical improvements, the mathematical foundations of phylogenetic compression remain little understood. General formulations of compression optimization are fundamentally NP-hard [12,13]. Yet, the simplified workflow of phylogenetic compression consistently achieves striking compression ratios. This paradox suggests that modern microbial collections have a highly pronounced combinatorial structure that phylogenetic compression exploits with unusual efficacy. However, a formal characterization of these underlying mechanics is still lacking.

Here, we introduce the first formal mathematical model of phylogenetic compression. We analyze a specific setting in which genome collections are represented as binary matrices – including SNP, *k*-mer, unitig, and unique-row matrices – and compressed with Run-Length Encoding (RLE). We prove that without structural assumptions, finding an optimal column ordering to minimize RLE size is NP-hard. However, when the genomes follow the Infinite Sites Model (ISM), Neighbor Joining (NJ) yields an optimal ordering in polynomial time. Finally, we experimentally validate these theoretical predictions on real bacterial datasets using an exact Traveling Salesperson Problem (TSP) solver. We demonstrate that, despite biological and technical deviations from ISM, NJ orderings achieve near-optimal compression across different diversity levels, matrix representation, and *k*-mer ranges, with UPGMA performing comparably well.

## 2 Methods

### 2.1 RLE compression of SNP, *k*-mer, unitig, and unique-row matrices

Let us have a bacterial genome collection *C* = {*G*_1_, …, *G*_*n*_}, we consider a particular form of the compression problem. We assume that the genome collection is represented as a binary matrix (e.g., of *k*-mer presence/absence), where each column corresponds to an individual genome and each row to a binary attribute; such representations are increasingly popular across bioinformatics, including SNP matrices, *k*-mer matrices, unitig matrices, and unique-row matrices, and underly many current indexing methods [10,14].

We assume that the matrix-represented collection is to be compressed by Run-Length Encoding (RLE) as a low-level compression method; this measures compressed size as the total number of runs across all rows, with each row processed independently. This framework allows us to study the column-reordering step as an optimization problem and characterize the conditions under which phylogeny-guided ordering is provably optimal.

### 2.2 Optimal column ordering under RLE compression is NP-hard for arbitrary data

Given a binary matrix, the objective of finding a column reordering minimizing the total RLE compression size is called *RLE binary matrix compression* (RBMC) problem.

#### Problem 1 (RBMC).

***Input***: *Binary matrix A* ∈ {0, 1}^*m×n*^

***Output***: *Column permutation of A that minimizes the total number of row-wise runs*

When no assumptions are made about its structure, the problem is NP-hard. In short, for any binary matrix, we can construct a complete graph *G* = (*V, E*) where each vertex *v*_*i*_ *∈ V* is the *i*-th column of *A*, and each edge (*v*_*i*_, *v*_*j*_) *∈ E* is weighted by the Hamming distance of the associated columns. Minimizing the total run count is then equivalent to finding the minimum-weight open Hamiltonian path in *G*, an open-path variant of the Traveling Salesperson Problem (TSP), which gives the NP-hardness (**Note S1**).

#### Theorem 1.

*The RBMC problem is NP-hard*.

The reduction gives an algorithm for computing the optimal RBMC orders using TSP solvers. First, we compute the pairwise column Hamming distances, then formulate the corresponding TSP instance, and finally solve it using a TSP solver such as Concorde [15]. With a small modification in the problem formulation, we can also obtain the worst-case order (**Note S2**).

### 2.3 The Infinite Sites Model (ISM) enables combinatorial modeling of genomes

Binary matrices that represent bacterial collections exhibit profound internal structure, enabling us to impose assumptions about the genomes via evolutionary modeling.

We adopt the classical *perfect phylogeny model* [16] (**Figure 1**a). A perfect phylogeny is a rooted tree in which the root represents an ancestral node with no derived characters, and each character appears exactly once and is never lost. When characters are binary, the phylogeny can be represented as a binary matrix *M* whose rows correspond to characters and columns correspond to objects (e.g., genomes); the value *M*_*ij*_ = 1 indicates that object *j* carries character *i*. Such a matrix is called a *perfect phylogeny matrix* (PPM). Perfect phylogenies have been widely studied [17,18,19], including efficient reconstruction algorithms from a PPM [16]. A central result for our work is the necessary and sufficient condition for a given binary matrix to be a PPM.

**Fig. 1:**
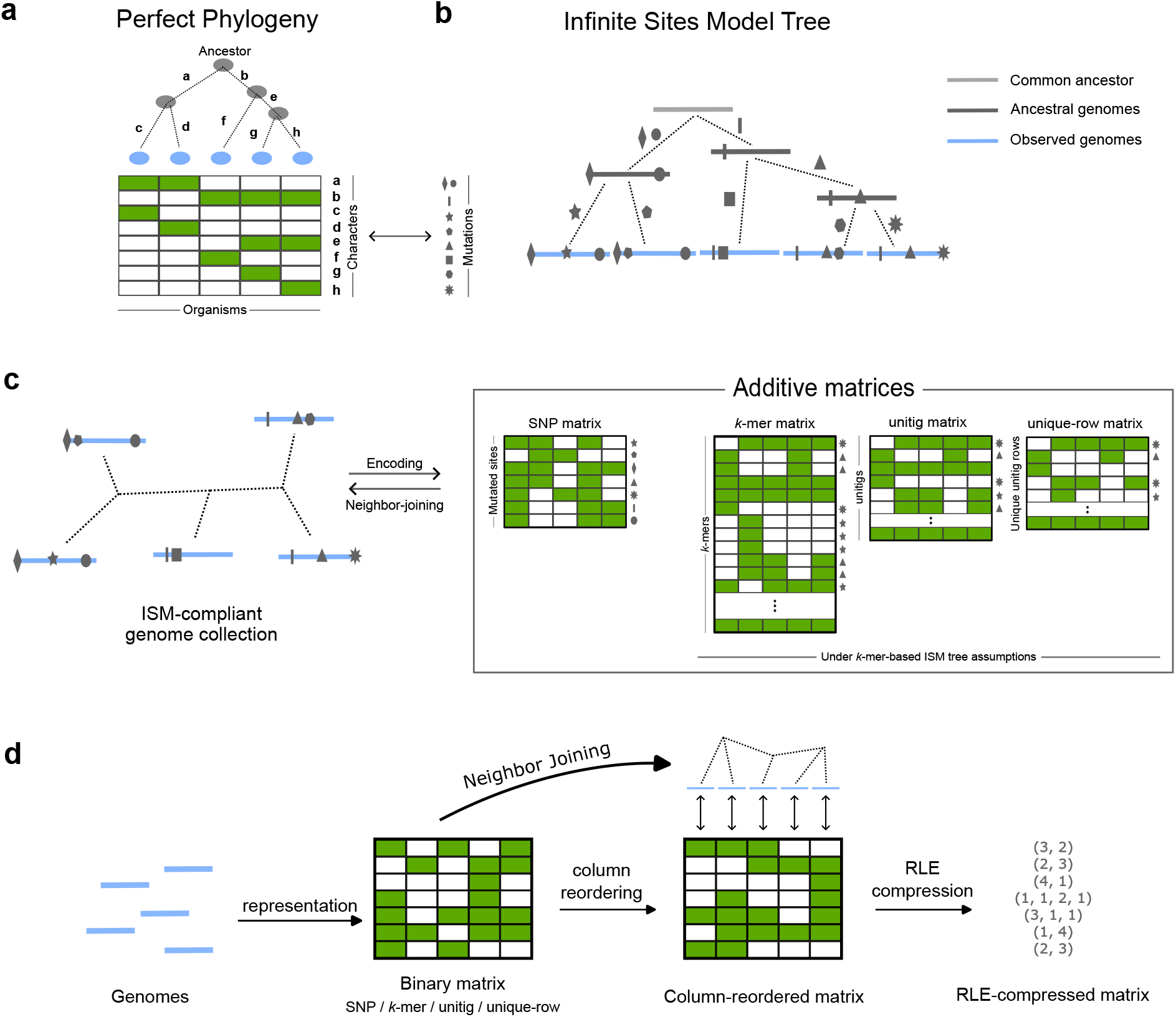
Overview of our methodology. **(a) Perfect phylogeny:** a classical tree-based model of character emergence. **(b) The Infinite Sites Model (ISM)** instantiates a perfect phylogeny for genomes and point mutations, producing a rooted binary tree. **(c) Binary matrices representing genome collections:** SNP, *k*-mer, unitig and unique-row matrices derived from ISM-compliant genomes satisfy the four-gamete condition and are thus ISM-compliant, inheriting the additive structure between columns which can be recovered via Neighbor Joining. **(d) A simplified protocol of phylogenetic compression:** for ISM-compliant matrices, phylogeny-guided compression performs optimally under the ISM assumptions.

#### Lemma 1

*([20,21]). Let M be a binary matrix. For each row k, let T*_*k*_ *be the set of columns where row k contains a* 1. *Then M is a PPM if and only if for every pair of rows i, j, the sets T*_*i*_ *and T*_*j*_ *are either disjoint or one contains the other*.

Perfect phylogenies naturally arise in bacterial genomics under the *Infinite Sites Model* (ISM) [22,23], which assumes that each genomic position mutates at most once. This simplified yet highly tractable model represents genomes as long strings, derived from a mutation-free ancestor through successive mutations at novel positions. More specifically, the ISM assumes that (1) each new mutation arises at a unique genomic position (no recurrence or reversal), and (2) there is no recombination. These assumptions enforce purely vertical inheritance: each position mutates at most once in a rooted tree, thereby delineating a unique clade and implying a perfect phylogeny; furthermore, the tree is without polytomies.

While the ISM remains a cornerstone for modeling genetic drift and mutation dynamics, our interest lies in the combinatorial patterns it guarantees; we formalize this combinatorial structure as a so-called *ISM tree* (**Figure 1b**).

#### Definition 1 (ISM tree).

*Let C* = {*G*_1_, …, *G*_*n*_} *be a genome collection composed of a set of equal length genomes. We call a rooted binary tree 𝒯 an ISM tree of the collection if*

1. *The root of 𝒯 contains no mutated positions*,
2. *the leaves of 𝒯 are exactly {G*_1_, …, *G*_*n*_},
3. *every edge is labeled by a set of mutations at unique positions, i*.*e*., *one position never appears in more than one edge label*,
4. *each descendant inherits the mutations accumulated on all edges from root to that node*,
5. *edge weights are derived as the number of mutations in the corresponding edge label. We call a collection C ISM-compliant if an ISM tree of C exists*.

### 2.4 ISM provides mathematical guarantees on the data structure of genome collections

The complete evolutionary history of real genome collections is rarely known, in particular the directionality. While mutations are usually easy to identify, for instance by comparison of all genomes to a fixed reference or via multiple sequence alignment, evolutionary directionality would require using an outgroup. This implies we need to work with unrooted trees.

This motivates a generalization of ISM trees to the unrooted case, reflecting that our reference genome may not be the true common ancestor. Given a binary matrix, we generalize the characterization of the perfect phylogeny matrix (**Lemma 1**), which relies on referencing the root, and introduce the concept of an *ISM-compliant matrix*.

#### Definition 2 (ISM-compliant matrix).

*Let M ∈* {0, 1}^*m×n*^ *be a binary matrix. For each row i, define two subsets of column indices T*_*i*_ = {*j* | *M*_*ij*_ = 1*} and F*_*i*_ = {*j* | *M*_*ij*_ = 0*}. Then M is called ISM-compliant if for each pair of rows r, s, at least one of these four intersections is empty T*_*r*_ ∩ *T*_*s*_, *T*_*r*_ ∩ *F*_*s*_, *F*_*r*_ ∩ *F*_*s*_, *F*_*r*_ ∩ *T*_*s*_.

ISM compliance of a matrix is equivalent to the *four-gamete condition* stating that no pair of rows contains all four binary patterns (00, 01, 10, 11), a criterion widely used for recombination detection [24] and perfect-phylogeny algorithms [25]. Naive ISM-checking requires 𝒪 (*nm*^2^) time, but specialized data structures allow improvements to 𝒪 (*nm*) [16]. As follows from **Lemma 1** and **Definition 2**, ISM-compliant matrices strictly generalize perfect phylogeny matrices (**Note S3**).

Every ISM-compliant matrix has an important structural property: pairwise Hamming distances between its columns are additive. A metric *d* on a finite set *S* is called *additive* if there exists a weighted tree *T* whose leaves are the elements of *S* such that for any *x, y*, the unique path-length in *T* connecting *x* and *y* is exactly *d*(*x, y*). ISM-compliant matrices inherit this property thanks to the four-gamete condition [26] (**Note S4**).

#### Theorem 2.

*Hamming distances induced by the columns of an ISM-compliant matrix are additive*.

When a genome collection is ISM-compliant, many of the commonly used representation matrices are ISM-compliant as well, although some require additional assumptions (**Figure 1c**). The SNP matrix is ISM-compliant under the condition of a compatible reference genome, i.e., a genome *X* such that it does not introduce a novel nucleotide in any position with respect to the collection (**Note S5**); the *k*-mer and unitig matrices are ISM-compliant under the assumptions of a *k*-mer-based ISM tree, as well as any matrix that is derived from those by deduplication of rows (**Notes S6–S7**).

#### Theorem 3.

*Let C* = {*G*_1_, …, *G*_*n*_*} be an ISM-compliant genome collection. Then:*

1. *The SNP matrix built on C with a compatible reference genome is ISM-compliant*.
2. *If C additionally forms the leaves of a k-mer-based ISM tree, then the k-mer matrix K, the unitig matrix U, and the unique-row matrix Q (created by deduplicating all rows of U) built on C are all ISM-compliant*.

### 2.5 ISM-compliant genome collections can be optimally reordered by Neighbor Joining

Altogether, when a binary matrix representing a genome collection is ISM-compliant, the RBMC problem simplifies dramatically. As the Hamming distances between columns become additive (**Theorem 2**), the distances can be perfectly explained by a tree, which breaks the NP-hardness. The central question is then whether the underlying tree can be reliably recovered. Although *some* tree can always be inferred by an arbitrary distance-based method, it may not explain the original distances.

It turns out the tree can be reconstructed robustly in 𝒪(*n*^3^) time by the Neighbor-Joining (NJ) algorithm. NJ provides the unrooted tree topology that is uniquely determined and corresponds (up to an isomorphism) to the ISM tree with its root removed, while placing the observed genomes as leaves, in a manner that exactly represents the observed mutational distances [27]. Moreover, the algorithm is robust: even for non-additive distances, it reconstructs a tree minimizing the balanced minimum evolution criterion [28]. An overview of the theoretical properties of NJ is given in **Note S8**.

#### Lemma 2

*([27], reformulated). Let T*_*u*_ *be an unrooted tree with shortest branch length s, and let 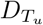 be the set of pairwise leaf distances of T*_*u*_. *Let D*_*m*_ *be a set of distances from* 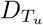, *modified by some amount. If all pairwise distances in D*_*m*_ *deviate from their true values in* 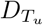 *by less than s/*2, *then the NJ algorithm reconstructs T*_*u*_ *exactly. In particular, if the distances are additive, exact reconstruction is guaranteed*.

Once the tree is known, finding an optimal column permutation becomes equivalent to identifying the shortest open Hamiltonian path in that tree. Such a path can be obtained by a simple depth-first traversal, with the only remaining degree of freedom being the choice of the two endpoint leaves that minimize the total path length.

This perfectly reflects how phylogenetic compression can proceed for such representations (**Figure 1d**): first, encode our data modeled by the ISM as an ISM-compliant matrix (e.g., *k*-mer matrix on data from a *k*-mer-based ISM tree); then, infer the phylogeny of the collection from the binary vectors using NJ; and lastly, reorder the columns of the matrix according to the phylogenetic tree left-to-right before applying the RLE compression. Under the assumption stated above, phylogenetic compression then yields optimal compression for these four common matrix representations of genome collections (**Note S9**).

#### Theorem 4.

*Let C* = {*G*_1_, …, *G*_*n*_*}be an ISM-compliant genome collection forming the leaves of a k-mer-based ISM tree, and let M* ^(*S*)^, *M* ^(*K*)^, *M* ^(*U*)^, *M* ^(*Q*)^ *be the associated SNP (with a compatible reference genome), k-mer, unitig, and unique-row matrices, respectively*.

*Then all four matrices are ISM-compliant. Moreover, NJ applied to the Hamming distances between their columns recovers the same unrooted tree topology 𝒯* . *An optimal solution to the RBMC problem can be obtained in 𝒪* (*n*^3^) *time by choosing branch rotations so that two leaves at maximum distance in 𝒯 are placed at opposite ends. Furthermore, any left-to-right leaf order of 𝒯 is optimal up to an additive term bounded by the diameter of 𝒯* .

## 3 Experimental evaluation

At first glance, we might think that real bacterial datasets are nowhere near the assumptions of our ISM. Genomes are finite and vary in length; individual nucleotide positions may mutate repeatedly or revert; genomes routinely undergo recombination and horizontal gene transfer; and genomic variation includes not only single nucleotide variants, but also small- and large-scale insertions, deletions, and structural rearrangements. These violations are even more pronounced at the *k*-mer level, where a single *k*-mer may occur multiple times within the same genome and be affected by multiple nearby mutations.

Yet our experiments on real bacterial datasets – ordered phylogenetically or according to exact solutions from a TSP solver – showed a different picture: a surprising agreement with the theoretical model.

### 3.1 NJ-orderings provide near-optimal compression across dataset diversities

We first asked how well TSP vs. NJ compress on datasets of varying diversity. We implemented a Snakemake pipeline (**Note S10**) to measure the total RLE runs of *k*-mer, unitig, and unique-row, based on random, optimal, worst-case and NJ-guided column orderings. We used it to analyze a benchmark of three datasets reflecting different RLE compression regimes (**Note S11**): 1) `ngono`, a single-species dataset that is easily compressible; 2) `ngono-spneumo`, a two-species mix, which introduces orderings very badly compressible with RLE (alternating species orders); and 3) `diverse`, a mix of 539 species which are generally little compressible. For each, the pipeline first extracted monochromatic unitigs using Fulgor [10], used them to compute the *k*-mer, unitig and unique-row binary matrices, computed pairwise column Hamming distances, formulated the corresponding TSP instances, and solved them exactly using Concorde [15] to obtain the most and least RLE-compressible orderings (**Note S10**). Finally, the pipeline infers an NJ phylogeny by Attotree [29] analogously to how they are inferred in MiniPhy [4], based on Mash [30] sketch distances as a proxy to *k*-mer distance and the QuickTree [31] implementation of NJ.

We found that NJ yields very similar RLE compression as exact solutions from a TSP (**Table 1**). With the single-species dataset, all phylogeny-based reorderings remained within 3% of the optimal and achieved roughly a 5*×* improvement over a randomized baseline on all three matrices (**Table 1a**). The worst-case ordering yielded around 120% of the randomized baseline, leaving the randomized baseline far closer to the worst-case scenario than the optimal (and phylogeny-informed) one.

**Table 1:**
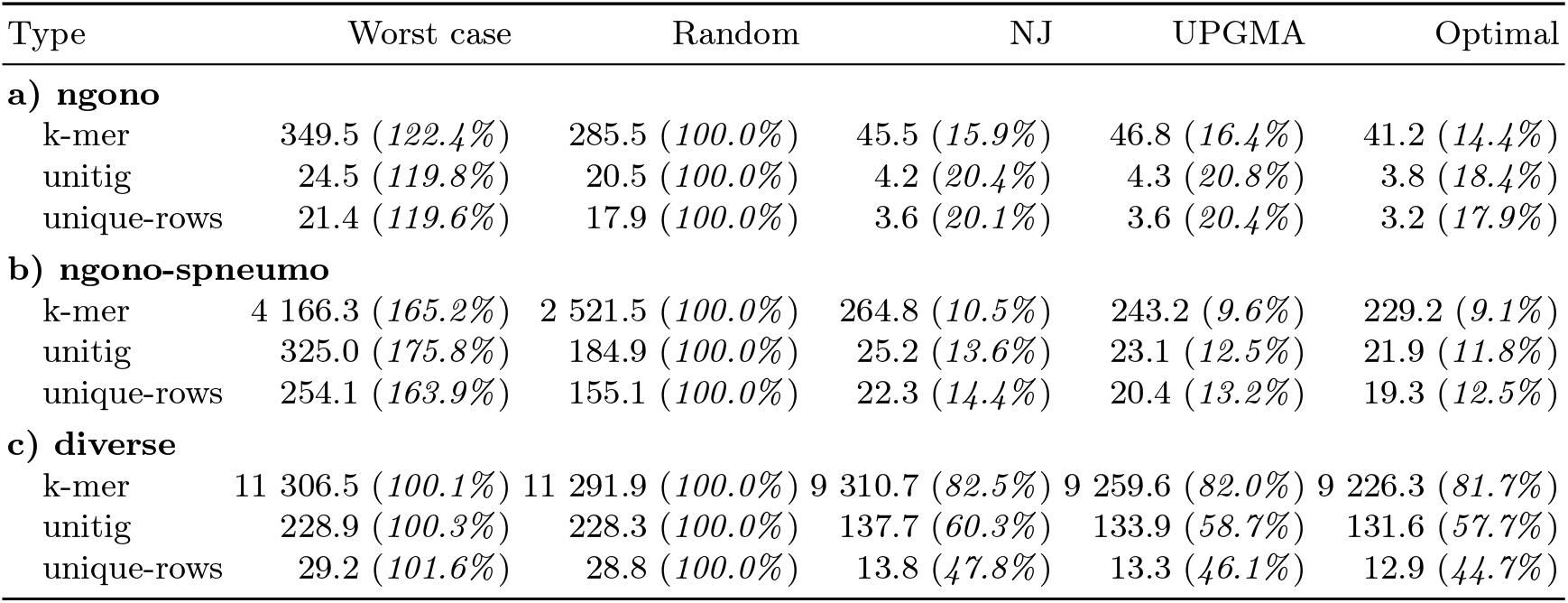
Effect of genome ordering on the RLE size of *k*-mer, unitig, and unique-row matrices across three datasets of increasing diversity. Each cell reports the number of runs (in millions) and, in parentheses, the relative size compared to the random-order baseline for the same dataset and matrix. Orderings considered are random, NJ, UPGMA, and the exact worst and best orderings obtained with a TSP solver. All results correspond to *N* = 1000 genomes and *k* = 31.

The two-species dataset showed qualitatively similar trends, but with substantially worse random and worst-case performance (**Table 1b**). The optimal solution improved randomized order by 10 fold and both NJ and UPGMA performed within 1% range from the optimal solution. The worst-case ordering reached over 160% of the randomized baseline and resulted from an order alternating the species’ genomes.

When the diversity grows (while preserving size), the ordering compression potential decreases (**Table 1c**). Here, the optimal solution resulted in only about 81% of randomized order on *k*-mer matrix, 57% on unitigs, and 44% on the unique rows. Notably, despite the signal being presumably dominated by horizontal gene transfer, the NJ order still approached the optimum with less than 1% difference. Interestingly, the absolute numbers of the runs in the unique-row matrix were much lower compared to *k*-mer or unitig matrices, a phenomenon likely caused by a large number of short unitigs shared by small amounts of genomes.

### 3.2 UPGMA yields compression comparable to NJ

While moving from the exponential worst-case complexity of TSP to *O*(*n*^3^), we asked whether it might be possible to use even less computationally demanding phylogenetic inference algorithm. As a natural candidate, we included also Unweighted Pair Group Method with Arithmetic Mean (UPGMA) [32] (directly supported by QuickTree and AttoTree); which additionally assumes ultrametric distances for correct reconstruction, but infers (rooted) trees in *O*(*n*^2^) with respect to the number of genomes.

Surprisingly, UPGMA yielded comparable RLE-compression performance to NJ (**Table 1**). For the single-species and two-species collections, the difference between NJ and UPGMA was within 1% relative to the randomized baseline. In the `ngono-spneumo` dataset, UPGMA produced marginally better compression than NJ, and in the `diverse` dataset it consistently outperformed NJ across all three matrix types. One possible explanation is that UPGMA’s simple averaging rule captures the local similarity structure that is most relevant for minimizing the number of runs. In other words, although UPGMA is not a biologically accurate model for our datasets, it may still well recover the important parts of the tree topology that matter most for compression.

### 3.3 The impact of reordering on RLE compression grows with the dataset size

We also examined the impact of the dataset size *n* on the improvement in compression with respect to the dataset diversity. We expected that larger single-species collections would yield greater improvements, while more diverse datasets would show more modest gains, with phylogeny-guided orderings remaining near-optimal regardless of size and diversity. As described in **Note S11**, for *k*-mer and unitig matrices the pipeline computed Hamming distances once in the matrix of the whole dataset, and nested subset runs were evaluated by extracting the corresponding columns to obtain the optimal, worst-case, and phylogeny-guided tours. The height of the *k*-mer and unitig matrix was therefore constant across all experiments. In the case of the unique-row matrix, the matrix was recomputed for each subsample before applying the rest of the pipeline.

Across all datasets and sample sizes (**Figure 2**), NJ-guided orderings remain consistently close to the optimal solution, typically within a few percentage points. Compression improves steadily as *n* increases, reflecting an initial saturation phase, where longer runs can form only once most *k*-mers or unitigs become present in the dataset subset. In single-species and two-species collections this saturation occurs quickly, whereas in the `diverse` dataset it appears only after roughly 100 genomes. Worst-case orderings are typically close to the random baseline, except in the two-species dataset, where the worst-case arrangement resulting from alternating genomes from the two species produces substantially poorer compression, with the gap widening as *n* increases.

**Fig. 2:**
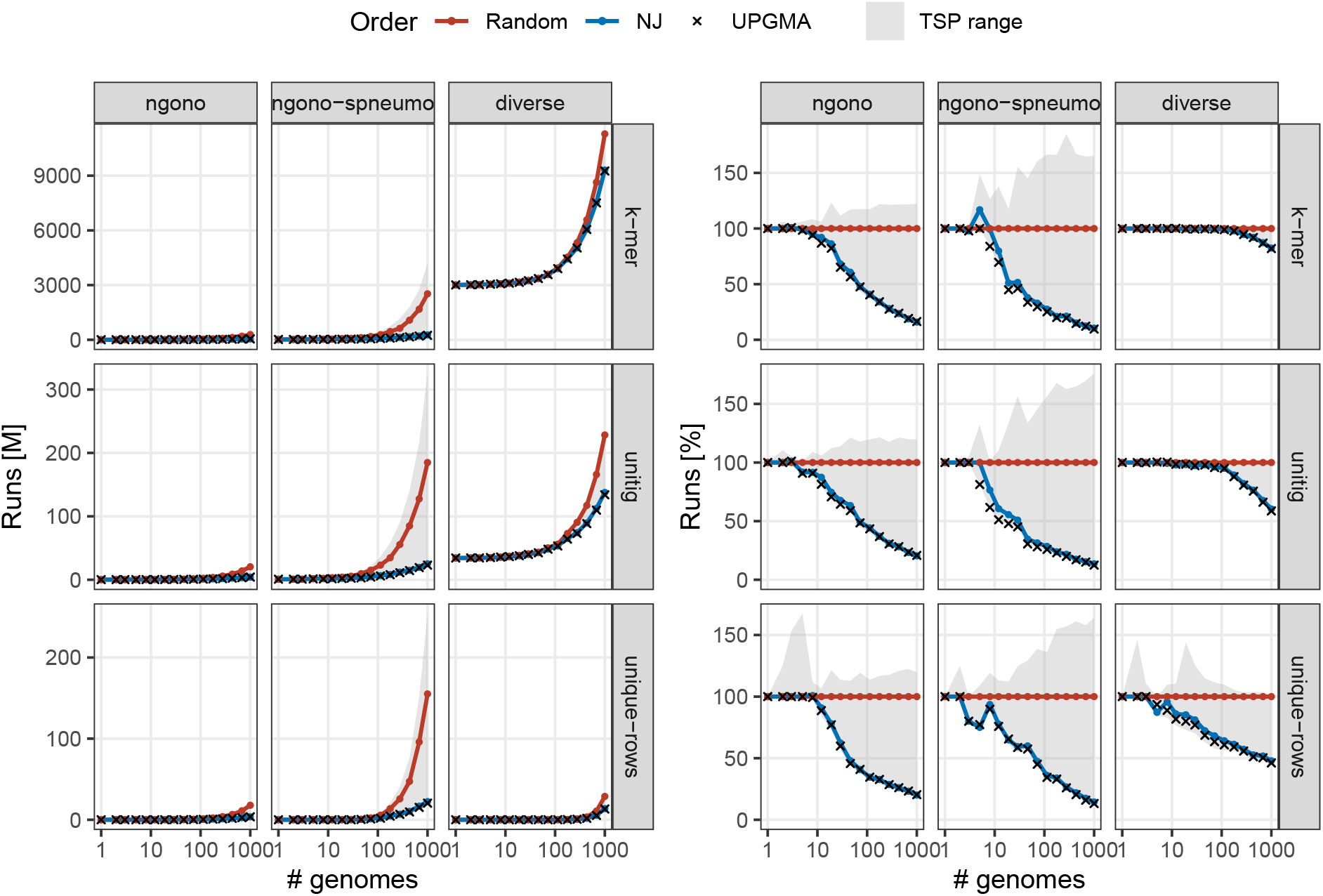
Impact of dataset size on compression performance. For each dataset, we evaluated the impact of increasing the number of genomes *n* on the row-wise run count in a RLE-compressed binary matrix. We generated nested subsamples of *n* genomes up to full dataset size and computed optimal, worst-case, random and phylogeny-guided orderings for each subset. The results show how compression improves with increasing sample size and how this behavior varies with dataset diversity.

The RLE compressibility of the *k*-mer, unitig, and unique-row matrices is generally similar. A notable exception occurs in the `diverse` dataset, where the unique-row matrix becomes more compressible earlier than the *k*-mer and unitig matrices. We conjecture that this difference is partly an artifact of the experimental design: because the unique-row matrix is rebuilt for each subsample, its structure reflects only the selected genomes, while the *k*-mer and unitig matrices include empty rows corresponding to *k*-mers or unitigs absent from the subset. On the other hand, these empty rows contribute only a constant baseline number of runs that is present across all subsets, and other types of runs that do not include this constant showed similar trends across matrix types (**Note S13**).

### 3.4 The results are robust to *k*-mer size

Finally, we evaluated the robustness to *k*-mer size, and found NJ still stays consistently near-optimal (**Figure 3**). Although the choice of *k* has a substantial impact on the size of the different matrix types and consequently their compressibility, the results further confirm the robustness of phylogeny-guided ordering improvements. Across all datasets, matrix types, and *k* values, both NJ and UPGMA orderings remain extremely close to the optimal TSP solution. Even in cases where the matrices are dense or significantly violate ISM assumptions, they consistently approach the optimum within a few percentage points.

**Fig. 3:**
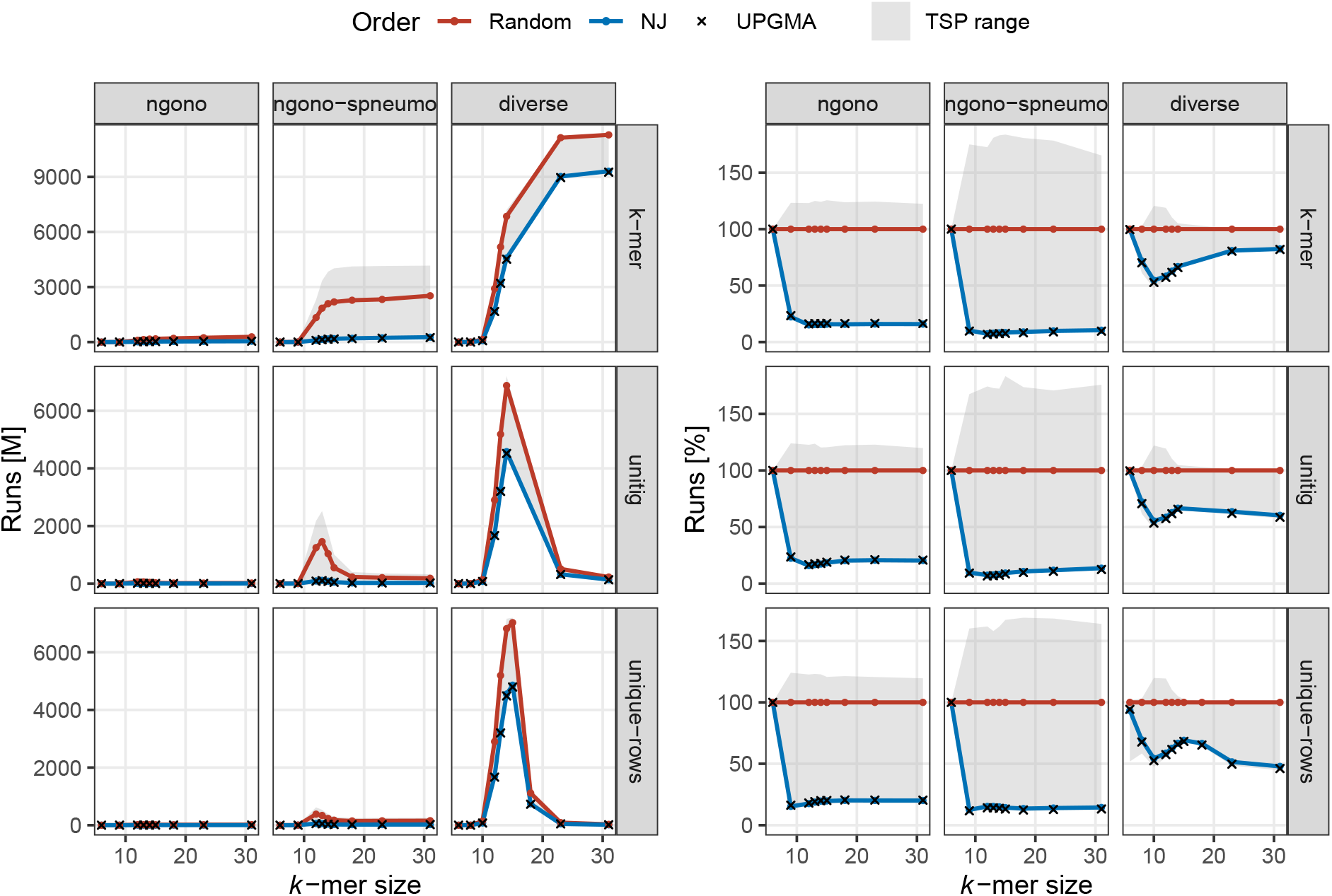
Effect of *k*-mer size on RLE compression performance. We evaluated how varying *k*-mer size *k* affects the number of runs in RLE-compressed binary matrices across three datasets (*N* = 1000 genomes) and three matrix representations. For each *k*, the figure shows the run count for random (red), NJ (green), and UPGMA (blue) column orderings, compared against the range bounded by the optimal and worst-case orderings obtained from TSP (gray). While the individual representations were considered with varying *k*, the trees were fixed and built with the default AttoTree *k*-mer size (*k* = 21).

The three matrix types exhibit distinct behaviors as *k* increases. The *k*-mer matrices begin in a highly dense regime for small *k*, since short *k*-mers appear in almost all genomes, and their height grows rapidly until stabilizing around *k* = 18. The unitig matrices show an even sharper initial increase, where for small *k*, the de Bruijn graph is highly fragmented, producing many short unitigs, which inflate the matrix height. As *k* increases, these fragments merge into longer unitigs, and the total number of unitigs decreases again around *k* = 18. The unique-row matrices follow a similar pattern but with substantially smaller absolute sizes, peaking around *k* = 15. Despite these differences in absolute matrix size, the relative compression performance measured against the randomized baseline shows much more stable behavior. After the dense, low-specificity regime at small *k*, the relative run counts settle quickly and remain nearly constant across the remaining *k* values, with only minor fluctuations in the diverse dataset. This suggests that once *k* is large enough for the matrices to reflect meaningful evolutionary structure, the relative advantage of different orderings becomes largely independent of the exact choice of *k*.

## 4 Discussion and conclusions

In this paper, we introduced and formally characterized a framework for modeling phylogenetic compression. Within this framework, we proved that the simplified phylogenetic protocol (constructing a tree from pairwise distances and reordering data in left-to-right leaf order) achieves the optimal RLE compression. We further showed that the compression problem itself can be expressed as a discrete optimization task equivalent to solving the Traveling Salesperson Problem (TSP) under Hamming distances, establishing that the general problem is NP-hard. Moreover, we defined the class of ISM-compliant matrices – a generalization of the binary matrices derived from data following the Infinite Sites Model (ISM) – that captures the essential structural properties required for phylogeny-based ordering to be optimal, and we demonstrated that several frequently used types of bioinformatics matrices belong to this class.

The empirical results demonstrate that many of the theoretical insights derived under ISM-like assumptions remain remarkably robust in practice. Even though real bacterial genomes violate the idealized model through homoplasy, recombination, and structural variation, their binary matrices still exhibit strong tree-like structure. As a consequence, phylogeny-based orderings consistently achieve compression close to the optimal TSP solution. In particular, the Neighbor-Joining algorithm (NJ) performed near-optimally not only for single-species datasets but also for more diverse collections across all our experiments. These findings suggest that bacterial genome space retains enough additive signal for simple distance-based methods to approximate the optimal ordering extremely well.

The main limitation of our experimental validation is the dataset size. To make exact TSP solutions tractable, we restricted our analyses to an upper bound of a limit of *N* = 1000 genomes to avoid scalability bottlenecks on our computational infrastructure; however, this size already corresponds to the scale of a typical batch size in phylogenetic compression. An additional limitation is the reliance on RLE as the low-level compression model, a simplification adopted for analytical tractability and ease of implementation. A more realistic model would incorporate also complex dictionary compressors, such as Lempel–Ziv variants [33].

This work brings several open questions. First, an important direction for future research is incorporating clustering into the theoretical model. In practice, phylogenetic compression operates on batches of phylogenetically related genomes, whereas our current model assumes a single tree constructed for the entire dataset. In principle, clustering could be integrated into our workflow by introducing additional dummy cites into the TSP formulation, reducing the optimization to the Clustered Traveling Salesperson Problem (CTSP) [34,35], which allows the solver to partition genomes into clusters. However, the resulting challenge is that individual matrices for each cluster would have different heights, which would need to be incorporated into the model.

Another interesting extension is the incorporation of vertical compression. While this study evaluated only row-wise RLE, the vertical direction presents substantial compression potential. Modern indexing methodologies implement such vertical partitioning and compression [36,37]. When advanced dictionary compressors such as XZ are applied to binary matrices, they successfully identify and exploit shared phrases across different rows. Although we partially capture this via evaluating the unique-row representations, this has been an incomplete solution. In fact, every split within an ISM tree segregates both genomes and variation, suggesting that the underlying principles developed for horizontal reordering are mathematically applicable to vertical compression as well.

Finally, an interesting question requiring additional formalization effort is the applicability of these principles to probabilistic data structures, such as Bloom filters. These structures are frequently used for large-scale indexing in practice [38,39] and can extensively benefit from compression. Prior implementations in MiniPhy and Phylign [4] successfully compressed Bloom filters with xz, though requiring an on-the-fly decompression during queries. The mathematical apparatus developed here may be adapted to Bloom filters by formally modeling hash collisions within the ISM framework.

Overall, this work provides both a theoretical foundation and an empirical validation for phylogenetic compression. By showing that simple, scalable methods can achieve near-optimal results despite substantial deviations from ideal evolutionary assumptions, it strengthens the case for using evolutionary structure as a guiding principle in the design of compressed genomic data structures in the broader spirit of compressive genomics [40] and aligning with recent broader theoretical perspectives that model evolution as constrained diffusion [41]. Together, these findings offer the first mathematical explanation for the effectiveness of phylogeny-guided compression and indexing. They reveal how the evolutionary structure inherent in bacterial genomes can be leveraged to overcome computational hardness barriers. In the long term, these insights contribute to the broader goal of developing entropy-scaling algorithms [42] capable of sublinear search in massive genomic collections, providing conceptual foundations to guide the design of future data structures and algorithms.

## Acknowledgments

This research was supported by the French National Research Agency (ANR) under Grant ANR-24-CE45-1226 for the REALL project (KB), and by the Czech Technical University in Prague under SGS 161-1612303D000 (VH). We are grateful to Pavel Veselý and Léo Ackermann for their constructive comments.

## Code Availability

The experimental evaluation pipeline is available at https://github.com/vercah/ISM-supplement.

