## Supplementary notes for "Why phylogenies compress so well: combinatorial guarantees under the Infinite Sites Model"

### Supplementary material for "Why phylogenies compress so well: combinatorial guarantees under the Infinite Sites Model"

#### S1 The RBMC problem is NP-hard

**Problem 1 (RBMC).**
***Input:*** *Binary matrix* $A \in \{0,1\}^{m \times n}$
***Output:*** *Column permutation of* $A$ *that minimizes the total number of row-wise runs*

**Theorem 1.** *The RBMC problem is NP-hard.*

*Proof.* Given a binary matrix $A \in \{0,1\}^{m \times n}$, we construct a complete graph $G = (V, E)$ where each vertex $v_i \in V$ is the $i$-th column of $A$. For every pair $i \neq j$, the edge $(v_i, v_j) \in E$ is assigned a weight equal to the Hamming distance $w(v_i, v_j) = \mathrm{Ham}(v_i, v_j)$.

The row-wise run count in $A$ can be directly expressed as $\sum_{j=1}^{n-1} w(v_j, v_{j+1}) + m$, where $m$ is the height of $A$, which is an additive constant with respect to the column order. Therefore, finding the order of columns that minimizes the total run count is equivalent to finding the minimum-weight open path in $G$ that goes through all nodes (i.e., Hamiltonian path), which is an instance of the open-path Traveling Salesperson Problem. To show that this problem is NP-hard under Hamming-distance weights, we reduce from the corresponding cycle version, which was proven to be NP-complete as the Hamming Traveling Salesperson Problem (HTSP) in [1].

We want to show that we can use the solution of the path TSP to solve the cycle variant. If we were able to compute the path solution in polynomial time, it would contradict the NP-hardness of the cycle variant.

Consider an instance of this cycle TSP variant given by our set of binary vectors $\{v_1, \ldots, v_n\}$ forming the columns of the matrix $A$ and the graph $G$ weighted by their pairwise Hamming distances. As described above, the shortest Hamiltonian path in $G$ is exactly an ordering of the vectors that minimizes the total number of row-wise runs in $A$.

To connect the shortest-cycle and shortest-path variants, we use a distance-equalization argument. We fix one of the vectors, without loss of generality $v_1$, and modify its distances so that it has an equal distance to any other vector, while preserving the order of vertices in the shortest Hamiltonian path. Let $D = \max_{i \neq 1} \mathrm{Ham}(v_1, v_i)$ be the maximum Hamming distance between $v_1$ and any other vector. For each $v_i, i \neq 1$, we add a constant $L - \mathrm{Ham}(v_1, v_i)$ to each of its distances $\mathrm{Ham}(v_i, v_j), j \neq i \wedge j \neq 1$. For binary vectors, we can do so by appending the same fixed binary suffix of appropriate length to every vector. Because adding the same constant to all edges incident on a vertex uniformly increases every tour length by twice that constant, the city order of the optimal tour over all cities remains unchanged.

Once $v_1$ is equidistant from every other vertex, it can be removed without affecting the relative ordering of any optimal path on the remaining $n - 1$ vertices. Solving this modified instance as a path TSP yields a shortest path on the vertices $v_2, \ldots, v_n$. Reinserting $v_1$ between the endpoints of this path closes it into a valid HTSP cycle on all $n$ vertices. Thus, any HTSP instance reduces in polynomial time to the path version, proving this version NP-hard as well. Consequently, the RBMC problem is NP-hard.

#### S2 Weight modification providing the worst case of the RBMC problem

**Lemma S1.** *Let $A$ be a binary matrix and let $G$ be the complete graph on its columns with edge weights $w(v_i, v_j) = \mathrm{Ham}(v_i, v_j)$. Let $W_{\max} = \max_{i \neq j} w(v_i, v_j)$ and define modified weights $w_m(v_i, v_j) = W_{\max} - w(v_i, v_j)$. Then the worst-case RBMC solution is obtained by solving the path TSP under the modified weights $w_m$.*

*Proof.* Under the modified weights, for any open Hamiltonian path $P = (v_1, v_2, \ldots, v_n)$, its total modified weight is

$$\sum_{k=1}^{n-1} w_m(v_k, v_{k+1}) = \sum_{k=1}^{n-1} \bigl(W_{\max} - w(v_k, v_{k+1})\bigr) = (n-1)W_{\max} - \sum_{k=1}^{n-1} w(v_k, v_{k+1}) \,.$$

Since $(n-1)W_{\max}$ is a constant, minimizing this modified weight is equivalent to maximizing the sum of the original Hamming distances along the path. By the same argument as in the proof of **Supplement S1**, the sum of Hamming distances is the total number of runs in $A$ up to an additive constant. Hence, a shortest path under $w_m$ gives a column ordering of $A$ with maximum total run count.

#### S3 ISM-compliant matrices strictly generalize perfect phylogeny matrices

The following classical result characterizes perfect phylogeny matrices (PPM).

**Lemma 1 ([2,3]).** *Let $M$ be a binary matrix. For each row $k$, let $T_k$ be the set of columns where row $k$ contains a 1. Then $M$ is a PPM if and only if for every pair of rows $i, j$, the sets $T_i$ and $T_j$ are either disjoint or one contains the other.*

ISM-compliant matrices relax the constraint of evolutionary directionality: the referenced features are no longer required to be those of the root, but are compared only among the observed genomes.

**Definition 2 (ISM-compliant matrix).** *Let $M \in \{0,1\}^{m \times n}$ be a binary matrix. For each row $i$, define two subsets of column indices $T_i = \{j \mid M_{ij} = 1\}$ and $F_i = \{j \mid M_{ij} = 0\}$. Then $M$ is called ISM-compliant if for each pair of rows $r, s$, at least one of these four intersections is empty $T_r \cap T_s$, $T_r \cap F_s$, $F_r \cap F_s$, $F_r \cap T_s$.*

Using these two characterizations, we can show the following.

**Lemma S2.** *Let $\mathcal{M}_{PP}$ denote the class of perfect phylogeny matrices, and let $\mathcal{M}_{ISM}$ denote the class of ISM-compliant binary matrices. Then $\mathcal{M}_{PP} \subsetneq \mathcal{M}_{ISM}$.*

*Proof.* Let us use the notation of a given binary matrix $M \in \{0,1\}^{m \times n}$ with the sets $T_k = \{j \mid M_{kj} = 1\}$ and $F_k = \{j \mid M_{kj} = 0\}$. Then $M \in \mathcal{M}_{PP}$ if for every pair of rows $i, j$, either the $T$ sets are disjoint ($T_i \cap T_j = \emptyset$) or one is a subset of the other ($T_i \cap F_j = \emptyset$ or $T_j \cap F_i = \emptyset$). Clearly, this satisfies the definition of ISM-compliance. Furthermore, the set of matrices that do not comply with this condition but satisfy for every pair of rows $F_i \cap F_j = \emptyset$ is ISM-compliant but not a perfect phylogeny matrix.

### S4 Hamming distances of the columns of an ISM-compliant matrix are additive

**Theorem 2.** *Hamming distances induced by the columns of an ISM-compliant matrix are additive.*

*Proof.* Recall the notation from **Definition 2**: $M \in \{0,1\}^{m \times n}$, $T_i = \{j \mid M_{ij} = 1\}$ and $F_i = \{j \mid M_{ij} = 0\}$. $M$ is ISM-compliant, so for each pair of its rows $r, s$, at least one of the intersections $T_r \cap T_s$, $T_r \cap F_s$, $F_r \cap T_s$, $F_r \cap F_s$ is empty. Equivalently, no pair of binary features given by rows $r, s$ per column can produce all four pairs $(0,0), (0,1), (1,0), (1,1)$ (the four-gamete condition).

We want to show that the columns in $M$, corresponding to binary representations of genomes, can be organized as leaves of a binary tree. A classical result of Peter Buneman [4] shows exactly this; each row $r \in M$ induces a *split* (bipartition) $(T_r, F_r)$ of the columns. By [4], if the splits satisfy the four-gamete condition, there is a unique binary tree whose edges realize exactly those splits and whose leaves correspond to the set of columns. In particular, each split $(T_r, F_r)$ corresponds to a unique edge whose removal separates the leaves into $T_r$ and $F_r$.

The Hamming distance $d(x, y)$ between two columns $x, y \in M$ counts the number of rows in which their values differ. In the split-induced tree, the row $r$ (which corresponds to exactly one edge $e_r$) separates $x$ and $y$ exactly when the path from $x$ to $y$ crosses $e_r$. Let us give weight $w(r)$ to the edge $e_r$ if the split is duplicated $w(r)$ times in the rows of $M$. Then the Hamming distance $d(x, y)$ equals the length of the path between $x$ and $y$ in the tree and is therefore additive.

#### S5 ISM-compliance of SNP matrices

**Definition S1 (SNP matrix).** Let $C = \{G_1, \ldots, G_n\}$ be a collection of genomes of equal length $L$ and $X$ be a genome of the same length. The SNP matrix $S \in \{0,1\}^{L\times n}$ of the collection $C$ with respect to $X$ as a reference genome is defined as

$$S_{ij} = \begin{cases} 1 & \text{if } G_j[i] \neq X[i], \\ 0 & \text{otherwise.} \end{cases}$$

**Definition S2 (Compatible reference genome).** Let $C = \{G_1, \ldots, G_n\}$ be a collection of equal-length genomes and let $X$ be a genome of the same length such that for any position $j$, $X[j] \in \{G_i[j] \mid G_i \in C\}$, i.e., $X$ does not introduce a novel nucleotide at any position with respect to the collection. We call $X$ a *compatible reference genome* of the collection $C$.

**Theorem S1.** *Let $C = \{G_1, \ldots, G_n\}$ be an ISM-compliant genome collection of equal length genomes with an ISM tree, and $X$ be a genome compatible with this collection. Then the SNP matrix built on the collection $C$ with the reference $X$ is ISM-compliant.*

*Proof.* Let us call the corresponding SNP matrix $S \in \{0,1\}^{L\times n}$, where $L$ is the length of the genomes, and use the notation of the subsets of column indices $T_i = \{j \mid S_{ij} = 1\}$, $F_i = \{j \mid S_{ij} = 0\}$ for each row $i$ in the SNP matrix $S$. We want to show that for each pair of rows $r, s$, at least one of the four intersections $T_r \cap T_s, T_r \cap F_s, F_r \cap T_s, F_r \cap F_s$ is empty.

Any row that is all-ones or all-zeros automatically satisfies this condition, so we restrict attention to the remaining (nontrivial) rows. Rows in an SNP matrix represent individual sites that mutate along the edges of the ISM tree $\mathcal{T}$, so each row is naturally associated with an edge of $\mathcal{T}$. This edge divides $\mathcal{T}$ into two subtrees, whose leaf sets correspond to the index sets $T_r, F_r, T_s, F_s$.

Let us choose two nontrivial rows $r, s$ in $S$. The edge associated with $r$ divides the tree into subtrees with leaf sets corresponding to $T_r$ and $F_r$. There are four possible scenarios, based on whether the edge $s$ lies within the subtree with leaves $T_r$ or within the subtree with leaves $F_r$.

– If the edge associated with $s$ lies within $T_r$, then either
  - $F_s \subseteq T_r$, therefore $F_s \cap F_r = \emptyset$,
  - $T_s \subseteq T_r$, therefore $T_s \cap F_r = \emptyset$.

– If it lies within $F_r$, then either
  - $F_s \subseteq F_r$, therefore $F_s \cap T_r = \emptyset$,
  - $T_s \subseteq F_r$, therefore $T_s \cap T_r = \emptyset$.

In all cases, at least one intersection must be empty, therefore the matrix $S$ is ISM-compliant.

#### S6 ISM compliance of $k$-mer matrices

**Definition S3 ($k$-mer matrix).** Let $\{G_1, \ldots, G_n\}$ be a collection of genomes, and fix $k \in \mathbb{N}$. Define $\{k_1, \ldots, k_D\}$ as the set of all distinct $k$-mers that appear in at least one genome of the collection. The $k$-mer matrix $K \in \{0,1\}^{D \times n}$ is defined as

$$K_{ij} = \begin{cases} 1 & \text{if } k_i \text{ occurs in } G_j, \\ 0 & \text{otherwise.} \end{cases}$$

To preserve the fundamental ISM properties for $k$-mer-based representations, we need to restrict how closely individual mutations can appear within a genome given a fixed $k$, since we want to prevent $k$-mers that appeared from mutated positions to disappear due to another mutation, and also prohibit the duplication of $k$-mers in independent branches of the phylogenetic tree. We formalize these extended assumptions as follows.

**Definition S4 ($k$-mer-based ISM tree).** Let $C = \{G_1, \ldots, G_n\}$ be a genome collection composed of a set of equal length genomes. We call a rooted weighted binary tree $\mathcal{T}$ a *$k$-mer-based ISM tree* of the collection if

1. the root of $\mathcal{T}$ contains no mutated positions,
2. the leaves of $\mathcal{T}$ are exactly $\{G_1, \ldots, G_n\}$,
3. every edge is labeled by a set of mutations, and any two mutations in $\mathcal{T}$ are at least $2k$ positions apart,
4. each descendant inherits the mutations accumulated on all edges from the root to that node,
5. in each genome, any $k$-mer appears at most once,
6. with each mutation in a genome, $k$ novel $k$-mers appear in that genome, where novel means such $k$-mer that only appears in the descendants of the given genome,
7. edge weights are defined as the number of mutations in the corresponding edge label.

**Theorem S2.** *Let $C = \{G_1, \ldots, G_n\}$ be a genome collection of genomes forming the leaves of a $k$-mer-based ISM tree. Then the $k$-mer matrix $K$ built on the collection $C$ is ISM-compliant.*

*Proof.* Under the $k$-mer-based ISM tree assumptions, one (single-nucleotide) mutation causes exactly $k$ $k$-mers to emerge and exactly $k$ $k$-mers to disappear. Each nontrivial $k$-mer is incident with exactly one edge. That edge cuts the leaf set into two complementary subsets that either contain the $k$-mer ($T_i$) or not ($F_i$).

The argument is similar to the one in the SNP matrix proof (Note S5). Every nontrivial $k$-mer labels exactly one edge, which splits the tree into two subtrees, whose leaf sets are $T_i$ and $F_i$. For another $k$-mer, its edge similarly determines $T_j$ and $F_j$. That edge lies either within the $T_i$ side or within the $F_i$ side. There are four possible scenarios as explained in Note S5. In all cases, at least one intersection is empty, therefore the matrix is ISM-compliant.

#### S7 ISM-compliance of unitig and unique-row matrices

The class of ISM-compliant matrices is invariant to row deletions and duplications.

**Lemma S3.** *Let $M$ be an $m \times n$ ISM-compliant matrix, and let $M_I$ be its submatrix obtained by retaining only the rows indexed by $I \subseteq \{1, \ldots, m\}$. Then $M_I$ is also ISM-compliant.*

*Proof.* Any pair of rows of $M_I$ is also a pair of rows of $M$. Since $M$ is ISM-compliant, the four-gamete condition (i.e., at least one of the intersections $T_r \cap T_s$, $T_r \cap F_s$, $F_r \cap T_s$, $F_r \cap F_s$ is empty) holds for every pair of rows of $M$, and therefore also for every pair of rows of $M_I$.

**Lemma S3** guarantees ISM-compliance for two commonly used matrices derived from the $k$-mer matrix by retaining representative rows.

First, consider the *unitig matrix*. Each row of this matrix represents a monochromatic unitig (see the formalization, e.g., in [5]). Each row of this matrix is obtained by retaining one representative row of the $k$-mer matrix from a unitig whose constituent $k$-mers all induce the same row of the $k$-mer matrix. This construction is well defined because the choice of representative does not affect the resulting row.

Second, consider the *unique-row matrix*, which consists of the distinct rows of the $k$-mer matrix. It is obtained by retaining one representative row from each equivalence class of identical rows.

**Corollary S1.** *Let $C$ be a genome collection whose $k$-mer matrix is ISM-compliant. Then the unitig and the unique-row matrices derived from $C$ are also ISM-compliant.*

Moreover, **Lemma S3** implies that any subsample of $k$-mers from the dataset yields an ISM-compliant matrix. Any subset of $k$-mers can be therefore used to encode the ISM-compliant collection as a binary matrix without violating its ISM-compliance, making this applicable even for matrices obtained via $k$-mer sampling techniques.

#### S8 Neighbor-Joining algorithm (NJ)

Here we describe the NJ algorithm in detail and summarize some results of its mathematical analysis that have been established in prior work.

Given a set of objects and their pairwise distances, the algorithm iteratively connects pairs of neighboring leaves, creating subtrees until an unrooted tree emerges. The core of the method is therefore the formula that determines the pair of neighboring leaves to be connected by a parental node. It is not sufficient to pick the leaves with the smallest distance, as it is demonstrated in **Figure S1**. Instead, the neighboring leaves are determined as the pair with the smallest value of the criterion

$$D_{ij} = d_{ij} - (r_i + r_j)\,,$$
$$r_i = \frac{1}{|L| - 2} \sum_{k \in L} d_{ik}\,, \tag{1}$$

where $L$ is the set of leaves and $d_{ij}$ is the distance between the leaves $i$ and $j$. This formulation ([6]) is not the one originally formulated by Saitou and Nei, however, they were proven equivalent ([7]). Studier and Keppler also provided the proof of the connection between the criterion $D_{ij}$ and the leaves' neighborhood.

**Theorem S3 ([8], [6]).** *If $i$ and $j$ are leaves chosen so that $D_{ij}$ is minimal, then $i$ and $j$ are neighbors.*

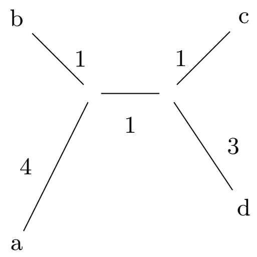

**Fig. S1:** Example of an unrooted tree where the pair of leaves connected by the shortest path ($b$ and $c$ with $d_{bc} = 3$) is not a neighboring one.

An iteration of the NJ algorithm runs as illustrated in **Figure S2**. Once the algorithm identifies a neighboring pair of leaves $i$ and $j$, it defines a parental node $k$ with the corresponding distances

$$d_{ik} = \frac{1}{2}(d_{ij} + r_i - r_j), d_{jk} = d_{ij} - d_{ik}\,, \tag{2}$$

and sets the new node's distances to other leaves to

$$d_{km} = \frac{1}{2}(d_{im} + d_{jm} - d_{ij})\,. \tag{3}$$

The leaves $i$ and $j$ are then removed from the list of leaves $L$ and $k$ is added, representing a new leaf for the next iteration. The whole process is described as **Algorithm 1**.

The algorithm performs well with both simulated and real data ([8], [10]), and the selection criterion $D_{ij}$ was proven to be unique and consistent among others that are linear, permutation equivariant, statistically consistent and based solely on distance ([11]). However, it is not immediately clear from **Equation (1)** what is the property that is being minimized by the NJ algorithm. It took more than ten years from the NJ introduction before the question was rigorously answered. Gascuel and Steel provide an insightful review of the subject ([7]).

The core of the answer lies in so called generalized Pauplin formula, which uses a weighted sum to obtain the total branch length in a tree.

**Definition S5 ([7], [12]).** Let $i$, $j$ be leaves in a weighted tree. Consider the path from $i$ to $j$ in the tree with $n$ interior nodes. Let $o_k$ denote the number of branches associated with the interior node $k$. The weight

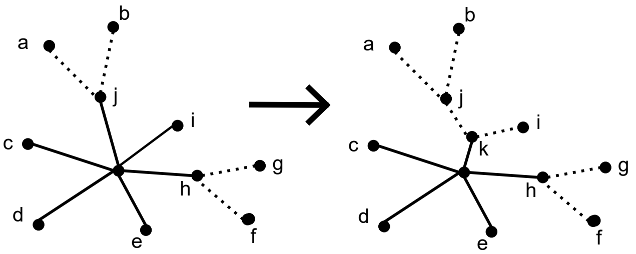

**Fig. S2:** One iteration of the NJ algorithm. Nodes $j$ and $i$ are identified as neighbors, and a parental node $k$ is created. Resolved nodes $i$ and $j$ are then removed from the set of current leaves and $k$ is added for the next iteration.

**Algorithm 1:** Neighbor-Joining (adopted from [9])

**Data:** set of leaves and their pairwise distances $d_{ij}$
**Result:** set of nodes and branch lengths of the corresponding unrooted tree
**Initialization:**
Define a set of leaves $L$ containing provided data;
Define a set of all nodes $T$, put $T = L$;
**while** *there are more than two leaves in* $L$ **do**
Determine the leaves $i$, $j$ for which $D_{ij}$ is minimal;
Define a new node $k$ and set $d_{km} = \frac{1}{2}(d_{im} + d_{jm} - d_{ij})$ for all $m \in L$;
Add $k$ to $T$ as parent of $i$, $j$ with edge lengths $d_{ik} = \frac{1}{2}(d_{ij} + r_i - r_j)$, $d_{jk} = d_{ij} - d_{ik}$;
Add $k$ to the set of leaves and remove $i$ and $j$;
**Termination:**
Connect the two remaining leaves $m$ and $n$ with an edge of length $d_{mn}$;

$w_{ij}$ is defined as

$$w_{ij} = \frac{1}{\prod_{k=1}^{n}(o_k - 1)} \,.$$

**Theorem S4 ([7], [12]).** *Let $l$ be the total sum of all branch lengths of a weighted tree. Then $l = \sum_{i,j} w_{ij} d_{ij}$.*

**Theorem S5 ([7], [13]).** *The NJ method, as defined by equations 1, 2, and 3 selects at each step as neighbors that pair of current leaves, which most decreases the whole tree length, as computed using the generalized Pauplin formula from* ***Theorem S4***.

#### S9 NJ solves the RBMC problem optimally for ISM-compliant matrices

**Theorem 4.** *Let $C = \{G_1, \ldots, G_n\}$ be an ISM-compliant genome collection forming the leaves of a $k$-mer-based ISM tree, and let $M^{(S)}, M^{(K)}, M^{(U)}, M^{(Q)}$ be the associated SNP (with a compatible reference genome), $k$-mer, unitig, and unique-row matrices, respectively.*

*Then all four matrices are ISM-compliant. Moreover, NJ applied to the Hamming distances between their columns recovers the same unrooted tree topology $\mathcal{T}$. An optimal solution to the RBMC problem can be obtained in $\mathcal{O}(n^3)$ time by choosing branch rotations so that two leaves at maximum distance in $\mathcal{T}$ are placed at opposite ends. Furthermore, any left-to-right leaf order of $\mathcal{T}$ is optimal up to an additive term bounded by the diameter of $\mathcal{T}$.*

*Proof. Part 1: Optimality of the leaf order.* Previous notes explain why all of the mentioned matrices are ISM-compliant. By **Theorem 2**, the Hamming distances between the columns of any ISM-compliant matrix $M$ are additive, therefore **Lemma 2** (from the main text) guarantees that the NJ algorithm produces a weighted unrooted tree $T$ whose leaf distances match the Hamming distances in $M$. By the proof of **Theorem 1**, finding a column permutation of $M$ that solves the **Problem 1** optimally is equivalent to finding a minimum-weight Hamiltonian path through the leaves of $T$. Since $T$ is the minimal spanning tree of all leaves, its shortest leaf traversal is realized by a standard depth-first traversal (DFT).

A DFT that starts at leaf $u$ and ends at leaf $v$ visits each edge twice, except those on the unique $u \to v$ path, which are traversed only once. If

$$W = \sum_{e \in E(T)} w(e)$$

is the total weight of $T$, then the length of the DFT route is

$$L(u, v) = 2W - d(u, v)$$

where $d(u, v)$ is the distance between $u$ and $v$ in $T$.

The DFT length can be further optimized by maximizing $d(u, v)$, i.e., by choosing the two leaves that are farthest apart as endpoints of the path. The best possible traversal length is therefore

$$L(u, v) = 2W - D,$$

where $D = \max_{\{u,v\}} d(u, v)$ is called the diameter of $T$.

*Part 2: Same tree topology for SNP, $k$-mer, unitig, and unique-row matrices.* Under the ISM, each mutation occurs exactly once on the underlying evolutionary tree. Consider any edge $e$ of this tree, and let $M(e)$ denote the number of mutations that occur along $e$. For any two genomes $G_i$ and $G_j$, their Hamming distance in each matrix type is the sum, over all edges on the path between $G_i$ and $G_j$, of a positive contribution per mutation on that edge. The magnitude of this contribution depends on the matrix type.

For the SNP matrix $M^{(S)}$, each mutation contributes exactly 1 to the Hamming distance between any pair of genomes whose path contains the corresponding edge. Thus the induced distances are additive with edge weights proportional to $M(e)$.

For the $k$-mer matrix $M^{(K)}$, by **Definition S4**, each point mutation creates exactly $k$ new $k$-mers and deletes $k$ old ones, contributing $2k$ to the Hamming distance for every pair of genomes separated by that mutation. Thus the induced distances are again additive, with edge weights equal to $2k\,M(e)$.

The unitig matrix $M^{(U)}$ is obtained from the $k$-mer matrix by merging certain rows that are identical across all genomes. As shown in **Lemma S3**, merging duplicated rows preserves additivity of the induced distances. Concretely, in the $k$-mer-based ISM tree, each edge $e$ corresponds to $2k\,M(e)$ rows that differ exactly on the genomes separated by $e$. When some of these rows are merged, the Hamming distances between genomes separated by $e$ decrease by the number of merged rows, which corresponds to reducing the weight of edge $e$ by the same amount. Since only duplicated rows are merged, the resulting edge weights remain strictly positive. Thus the unitig distances correspond to the same additive tree topology as the $k$-mer distances.

The unique-row matrix $M^{(Q)}$ is obtained from the unitig matrix by a further deduplication of identical rows. By the same argument via **Lemma S3**, this preserves additivity and positive edge weights, so the unique-row distances correspond to the same tree topology.

In all four cases, the pairwise Hamming distances are additive with respect to the same underlying tree topology, differing only by a scaling factor or by reductions in branch lengths that preserve positivity. It follows that NJ applied to $M^{(S)}$, $M^{(K)}$, $M^{(U)}$, or $M^{(Q)}$ reconstructs the same unrooted tree (up to branch rotations).

#### S10 Overview of the experimental pipeline

Our objective is to understand which ordering strategies produce the most compact RLE compression, and how these strategies behave across datasets, matrix types, and $k$-mer sizes. To answer these questions in a reproducible and scalable way, we implemented an evaluation pipeline using Snakemake [14]. The workflow coordinates all steps required to construct genome matrices, compute pairwise Hamming distances between columns, generate optimal and worst-case orderings via TSP formulations, infer phylogeny-based orderings, and finally quantify the resulting RLE-compressed sizes. The pipeline is modular and supports experiments across multiple datasets, $k$-mer sizes, and matrix types. A working example is available at https://github.com/vercah/ISM-supplement.

We conducted our experiments on several datasets of bacterial genomes obtained from public repositories of high-quality assemblies in FASTA format (**Table 1**). For each dataset, we first compiled a list of available assemblies and then extracted a random subset of 1000 genomes that served as the input of the pipeline. No additional preprocessing of the FASTA files was performed, as all subsequent steps of the pipeline operate directly on the assemblies.

To compute the unitigs of the collections, we used Fulgor [5], a software tool for indexing large collections of genomes by constructing a colored de Bruijn graph and compacting it into unitigs. Each genome corresponds to one color, and the resulting index can be exported into *dump files* containing the unitig sequences and their associated color sets.

We chose three types of binary matrices to represent the genome collections: the $k$-mer matrix, the unitig matrix, and the unique-row matrix. The $k$-mer matrix follows **Definition S3**, the unitig matrix follows the definition of monochromatic unitigs as defined in Fulgor [5]. In the unique-row matrix, each row corresponds to one color set, i.e., to a set of genomes sharing exactly the same unitigs. Theoretically, it can be derived from the unitig matrix by collapsing all identical rows. We showed that all three matrices are ISM-compliant when constructed from ISM-compliant data (**Theorem S2**, **Lemma S3**, **Corollary S1**). This motivates their use as test cases for evaluating the robustness of ISM-based predictions under realistic bacterial evolution.

To obtain column orderings that minimize or maximize the run-length-encoded size of the binary matrices, we applied the reduction of **Problem 1** to the TSP to be able to use an optimized TSP solver Concorde [15]. We used it also to compute the worst possible solution using distance modification from **Lemma S1**.

For phylogenetically guided column orderings, we inferred approximate phylogenetic trees for each dataset and extracted their left-to-right leaf orders. We employed two classical distance-based reconstruction methods implemented in Attotree [16]: Neighbor Joining (NJ), which is guaranteed to recover the correct topology for ISM-compliant matrices in our theoretical model (**Theorem 4**), and the Unweighted pair group method with arithmetic mean (UPGMA), which requires a stronger assumption of ultrametric distances to return the true tree. These two methods provided the phylogenetic heuristics for ordering matrix columns in our experiments.

In addition to phylogeny-based and TSP-based orderings, we included a randomized ordering, which served as a baseline model against which to compare the structured orderings.

To quantify how different column orderings affect the compressibility of the binary matrices, we measured the total number of row-wise runs. For a binary matrix $M \in \{0,1\}^{m \times n}$ and a fixed column ordering $\pi = (\pi_1, \ldots, \pi_n)$, the total number of runs is

$$\sum_{j=1}^{n-1} \mathrm{Ham}(M_{\bullet,\pi_j}, M_{\bullet,\pi_{j+1}}) + m,$$

where $m$ is the number of rows. The first term counts all bit changes across adjacent columns, and the additive constant accounts for the initial run in each row.

We also compared this row-wise measure with an alternative RLE strategy that linearizes the entire matrix and applies RLE globally. In this case, the total number of runs becomes

$$\sum_{j=1}^{n-1} \mathrm{Ham}(M_{\bullet,\pi_j}, M_{\bullet,\pi_{j+1}}) \ + \ \sum_{i=1}^{m-1} \mathrm{Ham}(M_{i,\pi_n}, M_{i+1,\pi_1}) \ + \ 1,$$

which removes the additive constant but introduces an additional term capturing transitions between the last column of one row and the first column of the next. Although we computed both quantities for all experiments, the differences between them were nearly constant and did not affect the qualitative conclusions. We therefore discuss only the row-wise run counts results in the experimental evaluation, which align with our theoretical model. A comparison of the two RLE strategies is shown in **Section S12**.

#### S11 Table of datasets

| Name | #genomes | #species | #distinct 31-mers | Description |
|---|---|---|---|---|
| ngono | 1000 | 1 | 4,117,063 | High-quality genomes of *Neisseria gonorrhoeae*. https://zenodo.org/records/15367750/files/part_54.tar |
| spneumo | 1000 | 1 | 16,268,711 | High-quality genomes of *Streptococcus pneumoniae*. https://zenodo.org/records/15367750/files/part_85.tar |
| ngono-spneumo | 1000 | 2 | 16,900,806 | A random subset of 500 *ngono* and 500 *spneumo* genomes. |
| diverse | 1000 | 539 | 3,009,067,743 | A random subset of 1000 genomes from all dust-bins of the phylogenetically compressed 661k collection. https://zenodo.org/records/15367750/ |
| ngono-rase | 1102 | 1 | 4,179,242 | Draft genome assemblies from Illumina HiSeq reads from the RASE database. https://github.com/karel-brinda/rase-db-ngonorrhoeae-gisp |

**Table 1:** Genome collections used in our experiments.

#### S12 Comparison of alternative RLE compression strategies

To evaluate whether our choice of RLE compression strategy influences the experimental results, we compared the row-wise approach used in our theoretical model (where each row begins a new run) with an alternative strategy that linearizes the entire matrix and applies RLE globally (**Figures S3 to S5**). Across all three main experiments, the relative differences between ordering methods remain highly consistent under both compression styles.

The only noticeable deviation appears in the `diverse` dataset. In the experiment varying dataset size, sparse rows introduce an additive constant that uniformly increases the total number of runs under the row-wise RLE strategy. In the experiment varying $k$-mer size, the difference between the two run types in the $k$-mer matrix grows with increasing $k$, reflecting the fact that the matrices become progressively sparser. This is the only setting in which the two strategies do not differ by a simple uniform shift. Nevertheless, the overall shape and behavior of the curves remain the same, and the gap would eventually converge to the height of the matrix once most $k$-mers become unique. For the remaining datasets, the differences between the two RLE strategies are negligible.

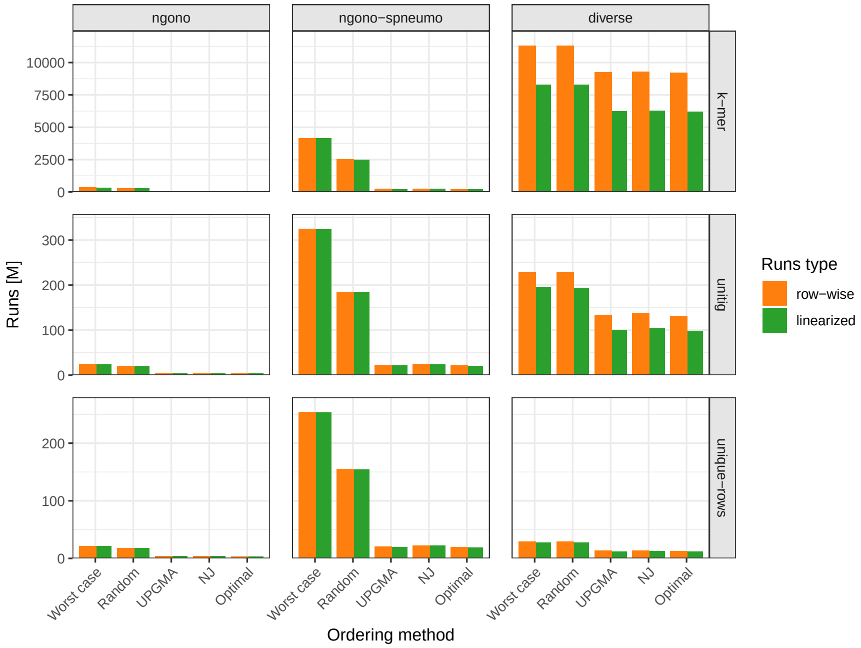

**Fig. S3: Comparison of RLE compression strategies across orderings and datasets.** Comparison of row-wise and linearized global RLE in the ordering experiment shows that relative differences between optimal, random, and phylogeny-guided orderings remain stable across both compression strategies.

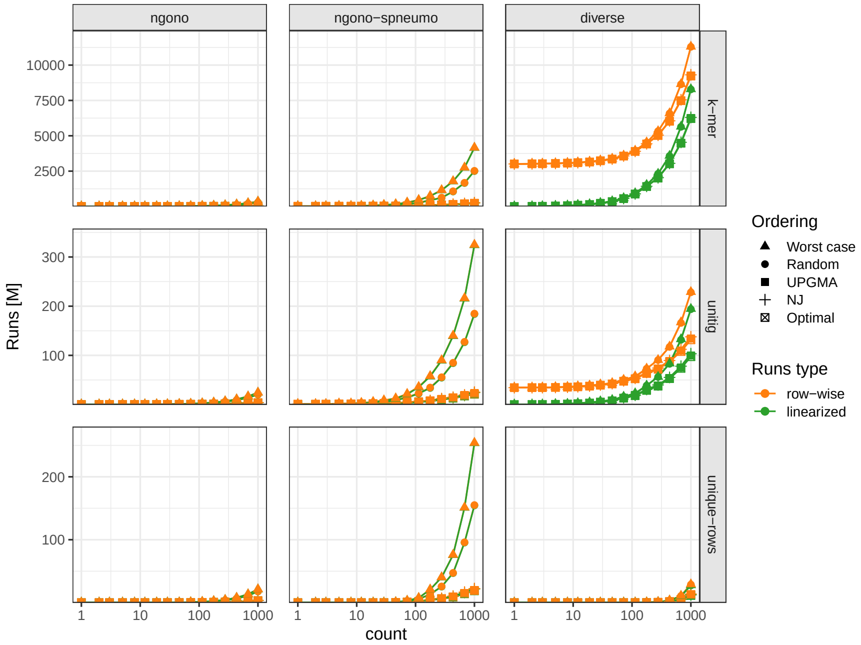

**Fig. S4: Comparison of RLE compression strategies across dataset sizes.** For increasing numbers of genomes, both RLE strategies yield nearly identical trends, with only a uniform shift in the `diverse` dataset due to sparse rows.

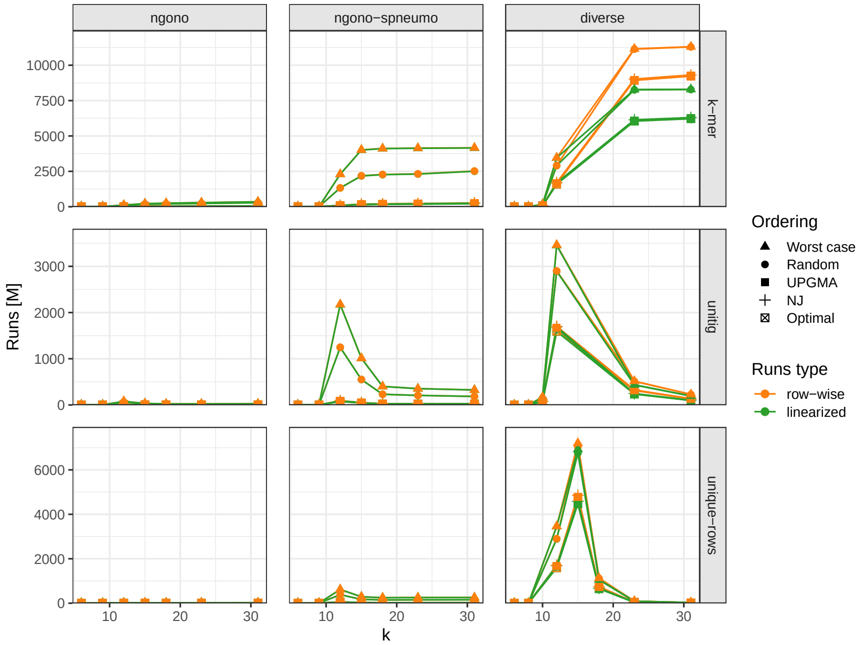

**Fig. S5: Comparison of RLE compression strategies across $k$-mer sizes.** Across all tested values of $k$, the two RLE approaches produce consistent relative ordering performance. The differences in the `diverse` dataset are caused by sparse rows, since many $k$-mers are unique.
